# HyDe: a Python Package for Genome-Scale Hybridization Detection

**DOI:** 10.1101/188037

**Authors:** Paul D. Blischak, Julia Chifman, Andrea D. Wolfe, Laura S. Kubatko

## Abstract

The analysis of hybridization and gene flow among closely related taxa is a common goal for researchers studying speciation and phylogeography. Many methods for hybridization detection use simple site pattern frequencies from observed genomic data and compare them to null models that predict an absence of gene flow. The theory underlying the detection of hybridization using these site pattern probabilities exploits the relationship between the coalescent process for gene trees within population trees and the process of mutation along the branches of the gene trees. For certain models, site patterns are predicted to occur in equal frequency (i.e., their difference is 0), producing a set of functions called *phylogenetic invariants*. In this paper we introduce HyDe, a software package for detecting hybridization using phylogenetic invariants arising under the coalescent model with hybridization. HyDe is written in Python, and can be used interactively or through the command line using pre-packaged scripts. We demonstrate the use of HyDe on simulated data, as well as on two empirical data sets from the literature. We focus in particular on identifying individual hybrids within population samples and on distinguishing between hybrid speciation and gene flow. HyDe is freely available as an open source Python package under the GNU GPL v3 on both GitHub (https://github.com/pblischak/HyDe) and the Python Package Index (PyPI: https://pypi.python.org/pypi/phyde).

---

It is increasingly recognized that a strictly bifurcating model of population descent is inadequate to describe the evolutionary history of many species, and is particularly common among plants, as well as some groups of animals (Mallet 2005; Baack and Rieseberg 2007; Mallet 2007). Hybridization and gene flow are processes that obscure this simple model, but separating the signal of admixture from other sources of incongruence, such as incomplete lineage sorting (ILS), is especially difficult (Maddison 1997). Early methods for detecting hybridization in the presence of ILS used estimated gene trees to detect deviations from the coalescent model under the expectation of a bifurcating species tree (Joly et al. 2009; Kubatko 2009; Meng and Kubatko 2009; Gerard et al. 2011). This work was later extended to include searches over network space to infer species networks with reticulate edges (Yu et al. 2011, 2012,2013, 2014; Solís-Lumis and Ané 2016). Because these network inference methods do not always scale to large numbers of species, other approaches for detecting hybridization that use genome-wide SNP data to test for admixture on rooted, four-or five-taxon trees have often been employed (“ABBA-BABA”-like methods; Green et al. 2010; Durand et al. 2011; Patterson et al. 2012; Martin et al. 2014; Eaton and Ree 2013; Pease and Hahn 2015). A common feature of these genome-wide methods for hybridization detection is their use of site patterns to test for deviations from the expected frequency under a neutral coalescent model with no introgression (Green et al. 2010; Durand et al. 2011).

The theoretical underpinnings of these site pattern-based inference methods originate from the study of invariants: functions of site pattern probabilities that contain information about the underlying relationships of the sampled species. For example, the D-statistic (Patterson et al. 2012) is based on invariants and uses the site patterns ABBA and BABA, which should occur in equal frequency in the absence of admixture. The earliest use of invariants to infer phylogenies was a pair of papers from Cavender and Felsenstein (1987) and Lake (1987), who separately derived functions to determine the correct topology for an unrooted quartet using binary and nucleotide data, respectively. Recent applications of phylogenetic invariants include proving identifiability (Chifman and Kubatko 2015), coalescent-based species tree inference (SVDquartets; Chifman and Kubatko 2014), sliding-window analyses of phylogenetic bipartitions (SplitSup; Allman et al. 2016), and the detection of hybridization (Green et al. 2010; Durand et al. 2011; Kubatko and Chifman 2015). The increased interest in methods using invariants is concomitant with the ability to collect genomic sequence data, allowing for accurate estimates of site pattern frequencies. Furthermore, because these methods are based on site pattern frequencies, they offer computationally tractable approaches for analyzing genome-scale data sets.

In this paper we introduce HyDe, a Python package for the detection of hybridization using phylogenetic invariants. HyDe automates the detection of hybridization across large numbers of species, and can conduct hypothesis tests at both the population and individual levels. A particular advantage of HyDe over other methods is its ability to assess individual-level variation in the amount of hybridization using per-individual hypothesis tests and individual bootstrap resampling in putative hybrid populations. The programming philosophy behind HyDe is to provide a low-level library of data structures and methods for computing site pattern probabilities and conducting hypothesis tests, along with a core set of Python scripts that use this library to automatically parse and analyze genomic data. This setup allows our software to be easily extended as new methods based on site pattern probabilities are developed. We describe the available features of the software in detail below and demonstrate the use of HyDe on both simulated and empirical data sets. To facilitate the use of HyDe for other researchers, we have provided all of the code for processing, plotting, and interpreting the results of these analyses.

## Description

### Model Background

In this section we provide a brief overview of the model used by HyDe for detecting hybridization. The theory behind the model, the derivation of the test statistics and corresponding asymptotic distributions, as well as assessments of statistical power and performance for different simulation settings can be found in Kubatko and Chifman (2015).

Consider a rooted, four-taxon network consisting of an outgroup and a triple of ingroup populations: two parental populations (P1 and P2) and a hybrid population (Hyb) that is a mixture of P1 and P2 [Figure 1]. Under this model, gene trees arise within the parental population trees following the coalescent process (Kingman 1982), where the hybrid population is either (1) sister to P1 with probability 1 - *γ* or (2) sister to P2 with probability *γ* (Meng and Kubatko 2009). Mutations are then placed on these gene trees according to the standard Markov models of nucleotide substitution [e.g., JC69 (Jukes and Cantor 1969), HKY85 (Hasegawa et al. 1985), GTR (Tavaré 1986)]. Given one sampled individual within each population, we can describe a probability distribution on the possible site patterns observed at the tips of the gene trees:

**Figure 1:**
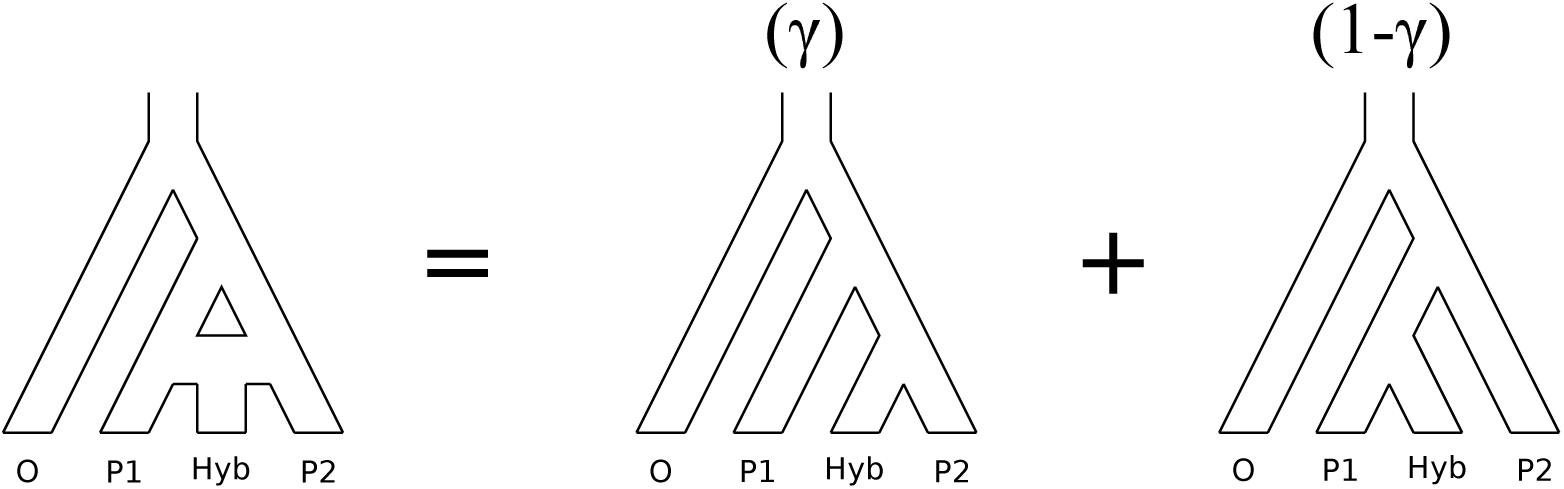
Illustration of the model for detecting hybridization using HyDe. The hybrid population (Hyb) is modeled as a mixture between two parental populations, P1 (1-*γ*) and P2 (*γ*).

*p*_*ijkl*_ = *P*(O = *i*, P1 = *j*, Hyb = *k*, P2 = *l*), with *i, j, k, l* ∈ {*A, G, C, T*}. Using nucleotide sequence data, we can also estimate these probabilities using the observed site patterns to get 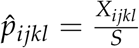, where *S* is the total number of observed sites and *X*_*ijkl*_ is the number of times that site pattern *ijkl* was observed. Implicit in this model is the assumption that each site evolves along its own gene tree, making unlinked sites the most appropriate input data. However, when the number of genes is large, it has been shown through simulations and empirical analyses that multilocus data provide a good approximation for the model and can be used to calculate site pattern probabilities (Chifman and Kubatko 2014; Tian and Kubatko 2017).

These site pattern probabilities form the basis of the test for hybridization that is implemented in HyDe, as well as several other methods. As we noted before, the D-statistic (Patterson et al. 2012) is based on invariants and uses the site pattern probabilities *p*_*ijji*_ and *p*_*ijij*_ to test for hybridization assuming a null model of no admixture:

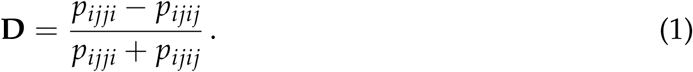

When admixture is absent, the expected value of **D** is 0, indicating that *p*_*ijji*_ and *p*_*ijij*_ should occur in equal frequency (i.e., they are invariant).

In a similar way, Kubatko and Chifman (2015) presented several different invariants-based test statistics that detect hybridization using linear phylogenetic invariants. Among the invariants that they tested, the ratio of *ƒ*_1_ = *p*_*iijj*_ - *p*_*ijij*_ and *ƒ*_2_ = *p*_*ijji*_ - *p*_*ijij*_ provided the most statistical power and is used to form the test statistic that is used by HyDe to detect hybridization. When the model holds, it can be shown that

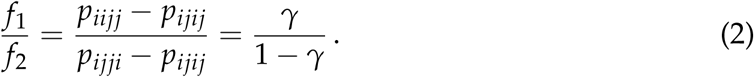

Deviation of the observed frequencies from this expectation is used to form a test statistic that asymptotically follows a standard Normal distribution, allowing formal hypothesis tests to be conducted (*H*_0_: *γ* = 0 vs. *H*_1_: *γ* > 0). Furthermore, because the ratio of *ƒ*_1_ and *ƒ*_2_ is a function of *γ*, we can estimate the amount of admixture directly from the observed site pattern frequencies:

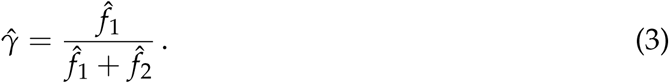

### Software Features

In this section we outline the main features of the HyDe software. Previously, Kubatko and Chifman (2015) used C code to test their new methods and distributed it in accordance with standard publishing practices. However, the code was only intended to evaluate performance for their method of hybridization detection, and not as a “distribution-level” piece of software. The new Python package that we present here serves that purpose, and provides additional functionality to accommodate more complex hybridization scenarios and sampling designs.

#### Multiple individuals per population

We have modified the calculation of the site pattern probabilities originally presented in Kubatko and Chifman (2015) to accommodate multiple individuals per population, rather than testing each sampled individual separately. We do this by considering all permutations of the individuals in the four populations involved in the hypothesis test. For example, if there are *N*_*O*_ individuals in the outgroup, *N*_*P*1_ individuals in parental population one, *N*_*Hyb*_ individuals in the putative hybrid population, and *N*_*P*2_ individuals in parental population two, then a total of *N*_*O*_ × *N*_*P*1_ × *N*_*Hyb*_ × *N*_*P*2_ quartets are used to calculate the site patterns for the hypothesis test. Including multiple individuals per population increases the sample size used to calculate the site pattern probabilities and can therefore lead to more accurate detection of hybridization.

#### Identifying individual hybrids

An underlying assumption of the coalescent with hybridization is that all the individuals in the hybrid population are admixed (i.e., hybrid speciation; Meng and Kubatko 2009). However, when gene flow, rather than hybrid speciation, is responsible for the introgression of genetic material into the admixed population, it is possible that not all individuals will contain these introgressed alleles. When hybridization detection is conducted on populations with a mix of hybrids and non-hybrids, it is possible for invariant-based tests to report significant results, even though the underlying assumption of uniform admixture is violated. To help with the detection of non-uniform introgression into the hybrid population, we have included two methods in HyDe that aim to detect variation in the amount of hybridization in the individuals sampled.

If not all of the individuals in a putative hybrid population are admixed, bootstrap resampling can reveal heterogeneity in the process of introgression when more or fewer hybrid individuals are included in each replicate. For example, if we are testing four diploid individuals in a putative hybrid and only two of them are 50:50 hybrids (*γ* = 0.5 for each), then the value of gamma for the whole population will be *γ* = 0.25. When we resample individuals with replacement and recalculate *γ*, we will get different values depending on how many times the hybrids are resampled (0 times: *γ* = 0.0; 1 time: *γ* = 0.125; 2 times: *γ* = 0.25; 3 times: *γ* = 0.375; 4 times: *γ* = 0.5). Because the process of hybridization is not uniform, the value of *γ* jumps between different values. This can also be visualized by plotting the distribution of *γ* across the bootstrap replicates (e.g., Figure 2).

**Figure 2:**
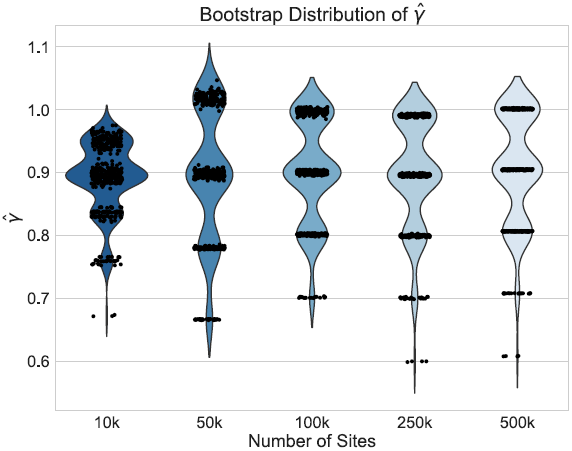
Violin plots of the distribution of 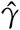 across 500 bootstrap replicates for 10 000, 50 000, 100 000, 250 000, and 500 000 sites for the non-uniform hybridization simulations. The black dots in each violin plot represent the actual values of 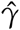 that were estimated by each replicate (black dots are jittered). The pattern of jumping between distinct values of *γ* with no estimates in between are a strong indication of non-uniform admixture.

We have also implemented methods that test all individuals in a putative hybrid separately while treating the parents and outgroup as populations. This approach allows hypothesis tests and estimates of *γ* to be calculated for every individual to see if it is a hybrid. A caveat with testing each individual is that the number of sites sampled must be enough to have statistical power to detect that hybridization has occurred (Kubatko and Chifman 2015).

#### Ambiguity codes and missing data

To allow more data to be used, and to account for the potentially large amounts of missing data that are common in high throughput sequencing data sets, we have implemented approaches that integrate over missing or ambiguous nucleotides by considering the possible resolutions of the observed bases into site patterns. As an example, consider the site pattern AGRG. There are two possible resolutions for the ambiguity code R: AGGG and AGAG. To account for this, we add 0.5 to the site pattern counts *X*_*ijjj*_ and *X*_*ijij*_. In general, for any site pattern containing ambiguous or missing bases (but not gaps), we find all possible resolutions and add one divided by the total number of resolutions to the corresponding site pattern counts.

### Software Design and Implementation

HyDe is implemented in Python and also uses Cython, a superset of the Python language that allows C/C++ capabilities to be incorporated into Python for better efficiency (Behnel et al. 2010). Installing HyDe requires several external Python modules [`numpy, cython, matplotlib, multiprocess, and seaborn`]. The goal behind our implementation of HyDe was to provide both pre-packaged scripts to conduct standard hybridization detection analyses, as well as a library of functions and data structures that researchers could use to customize their analyses or implement new tests based on site pattern probabilities. Below we describe the functionality of both of these interfaces, as well as the input files required to run HyDe.

#### Input files

The input files for running HyDe are plain text files containing the DNA sequence data and the map of individuals to populations. The DNA sequence data are expected to be in sequential Phylip format and the population map is a two-column table with individual names in the first column and the name of the population that it belongs to in the second column. The third input file that is required for individual-level testing and bootstrapping (optional for running a normal HyDe analysis) is a three column table of triples. A hypothesis test is then run for each triple using the first column as parent one, the second column as the putative hybrid, and the third column as parent two. The outgroup for each test is always the same and is specified separately at the command line. Output files from previous HyDe analyses can also be used to specify which triples are to be tested.

#### Command line interface

The command line interface for HyDe consists of a set of Python scripts that can be used to detect hybridization, filter results, conduct individual bootstrapping, and test for hybridization at the individual level. The three main scripts for detecting hybridization with HyDe provide the majority of the basic functionality that is needed to detect hybridization in an empirical data set and to assess if individuals in putative hybrids are all equally admixed. Each of these scripts also has a multithreaded version for parallelizing hypothesis tests across triples (in square brackets below).

- ***run_hyde.py*** [*run_hyde_mp.py*]: The *run_hyde.py* script detects hybridization by running a hypothesis test on all possible triples in a data set in all directions (i.e., a “full” HyDe analysis). There is also an option to supply a list of specific triples (three column text file) to test. The script outputs two files: one with all of the hypothesis tests that were conducted and one with only those hypothesis tests that were significant at an overall *α* = 0.05 level (after incorporating a Bonferonni correction) with estimates of *γ* between 0 and 1.
- ***individual_hyde.py*** [*individual_hyde_mp.py*]: The *individual_hyde.py* script runs separate hybridization detection analyses on each individual within a putative hybrid lineage. The only additional input needed for the script is a three column list of triples or a filtered results file from the *run_hyde.py* script.
- ***bootstrap_hyde.py*** [*bootstrap_hyde_mp.py*]: The *bootstrap_hyde.py* script performs bootstrap resampling of the individuals within a putative hybrid lineage and conducts a hypothesis test for each bootstrap replicate. Similar to the *individual_hyde.py* script, the script uses as input a table of triples or a filtered results file from a previous hybridization detection analysis.

The workflow that we envision for these scripts starts with using the *run_hyde.py* script to test for hybridization on all triples in all possible directions, and to filter out only those triples that have significant evidence for hybridization. Then, using the filtered output file, users can either test all of the individuals in each hybrid lineage or perform bootstrap resampling of the individuals using the *individual_hyde.py* and *bootstrap_hyde.py* scripts, respectively. Users with specific hypotheses that they want to test can simply create a text file listing the triples of interest to be run through any of the three scripts.

#### Python API

Each of the command line scripts described above makes use of an underlying Python library with built-in data structures for processing the hypothesis tests and bootstrapping analyses conducted by HyDe. This library is exposed through the Python module `phyde` (**P**ythonic **Hy**bridization **De**tection). The main data structures that are part of this module are listed below:

- ***HydeData***: The *HydeData* class is the primary data structure for reading in and storing DNA sequence data and maps assigning individuals to populations. This class also implements three of the main methods for detecting hybridization at the population level [`test_triple()`] and the individual level [`test_individuals()`], as well as bootstrapping individuals within populations [`bootstrap_triple()`] (see Box 1).
- ***HydeResult***: The *HydeResult* class parses the results file output by a hybridization detection analysis and stores the results as a Python dictionary (key-value pair). Values stored by the *HydeResult* class can be accessed by providing the name of the desired value and the names of the taxa in the triple of interest as arguments to the variable used when reading in the results. For example, if we read the results of a hybridization detection analysis into a variable named `results`, we would access the estimated value of 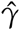 for the triple (`sp1, sp2, sp3`) using the following code: `results(“Gamma”, “sp1”, “sp2”, “sp3”)`.
- ***Bootstrap***: The *Bootstrap* class is similar to the *HydeResult* class except that it has built-in methods for parsing the format of the bootstrap output file written by the *bootstrap_hyde.py* script. Each bootstrap replicate is printed with a single line containing four pound symbols separating each tested triple (####\n). These results are parsed into a Python dictionary that can be used to access the bootstrap replicates for particular triples using the variable name, just like the *HydeResult* class [e.g., `bootstraps(“Gamma”, “sp1”, “sp2”, “sp3”)`].
- ***phyde.viz***: The *viz* submodule uses the `matplotlib` and `seaborn` libraries to implement basic functions for plotting the distribution of bootstrap replicates for any quantity calculated by the *bootstrap_hyde.py* script (Z-score, p-value, *γ*, etc.) for a specified triple. It does this by taking the name of a *Bootstrap* object variable, the name of the value to be plotted, and the names of the taxa in the triple of interest (Supplemental Materials: HyDe_SysBio.ipynb).

## Benchmarks

All code used to complete the simulations and example analyses are available as a Jupyter Notebook on Dryad (dryad.####). We have also included all output files needed to reproduce the tables and figures. Note that in this section we use the names `out, sp1, sp2`, and `sp3` for the names of the populations, rather than O, P1, Hyb, and P2. We do this to denote that these taxa have not yet been tested for hybridization, whereas the model is explicitly constructed to contain a hybrid species and its parents.

### Non-uniform Hybridization Simulations

To demonstrate the use of the methods in HyDe that are designed to identify individual hybrids, we set up a simulated example for four taxa (`out, sp1, sp2, sp3`) that intentionally violated the assumption of hybrid speciation by including a single hybrid individual in the population being tested for admixture (`sp2`). We simulated gene trees using the program `ms` (Hudson 2002) with population branch lengths of 1.0 coalescent unit assuming a model of coalescent independent sites (i.e., one gene tree per site; Kubatko and Chifman 2015) for 10 000, 50 000, 100 000, 250 000, and 500 000 sites. The parental populations were simulated with five individuals each and the outgroup had only one individual. The “admixed” population (`sp2`) had four non-admixed individuals that were most closely related to parental population `sp1`, and a single hybrid individual that was a 50:50 (*γ* = 0.5) mix between `sp1` and `sp3`. A single DNA base was simulated on each gene tree for the different numbers of sites using the program `Seq-Gen` (Rambaut and Grassly 1997). We rescaled gene tree branch lengths from coalescent units to mutation units using a population scaled mutation rate (*θ* = 4*N*_0_*μ*) of 0.01 per site and used a GTR+Gamma model of nucleotide substitution with three rate categories:

~~~
seq-gen-mGTR-s 0.01-l 1-r 1.0 0.2 10.0 0.75 3.2 1.6
-f 0.15 0.35 0.15 0.35-i 0.2-a 5.0-g 3
~~~

Output from `Seq-Gen` was formatted for input to HyDe using a Python script (*seqgen2hyde.py*; available on Dryad). Hybridization detection was completed for the simulated data sets with HyDe v0.4.1 using the *run_hyde.py, bootstrap_hyde.py* (500 bootstrap replicates), and *individual_hyde.py* scripts. Output files were processed and plotted in Python v2.7 using the `phyde, matplotlib`, and `seaborn` modules.

#### Results

Testing for hybridization at the population level using the *run_hyde.py* script produced a test statistic indicating that there was significant admixture in the `sp2` population at the *α* = 0.05 level when the number of sites was at least 50 000 (Table 1). The estimated values of 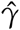 for these tests were close to 0.9, which is inconsistent with the data that we simulated. The value of *γ* that we would expect is 0.1. However, because this is an explicit violation of the coalescent model with hybridization, the formula for the estimation of *γ* is no longer valid.

**Table 1:** Results of the population-level hybridization detection analyses for the non-uniform hybridization simulation using the *run_hyde.py* script. Note that the result for the simulation with 10 000 sites was not significant at the *α* = 0.05 level (marked with _*_).

| # sites | Z-score | P-value | $\hat{\gamma}$ |
| --- | --- | --- | --- |
| 10 000* | 1.100 | 0.136 | 0.896 |
| 50 000 | 2.234 | 0.013 | 0.897 |
| 100 000 | 3.274 | $5.310 \times 10^{-4}$ | 0.900 |
| 250 000 | 5.420 | $2.973 \times 10^{-8}$ | 0.896 |
| 500 000 | 6.808 | $4.966 \times 10^{-12}$ | 0.905 |

If we did not already know that not all of the individuals were hybrids, we would mistakenly infer that the `sp2` population has ~90% of its genetic material inherited from population `sp3`. However, when we test each individual using the *individual_hyde.py* script, we correctly infer that only one of the individuals is admixed (individual ‘i10’, 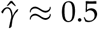) and that the rest of the individuals are not hybrids (Table 2). Bootstrap resampling of the individuals in population `sp2` also indicates that hybridization is not uniform. Figure 2 shows the distribution of 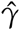 across all 500 bootstrap replicates and demonstrates that the value of 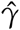 jumps between different values depending on the number of times the hybrid individual is resampled.

**Table 2:** Results of the individual-level hybridization detection analyses for the non-uniform hybridization simulation using the *individual_hyde.py* script. Individual ‘i10’ was the only hybrid in the population and was simulated with *γ* = 0.5.

| Individual | 10 000 |  |  | 50 000 |  |  | 100 000 |  |  | 250 000 |  |  | 500 000 |  |  |
| --- | --- | --- | --- | --- | --- | --- | --- | --- | --- | --- | --- | --- | --- | --- | --- |
| | Z-score | P-value | $\hat{\gamma}$ | Z-score | P-value | $\hat{\gamma}$ | Z-score | P-value | $\hat{\gamma}$ | Z-score | P-value | $\hat{\gamma}$ | Z-score | P-value | $\hat{\gamma}$ |
| i6 | 0.408 | 0.342 | n.s. | 0.054 | 0.478 | n.s. | 0.049 | 0.480 | n.s. | 0.387 | 0.349 | n.s. | -0.015 | 0.506 | n.s. |
| i7 | 0.241 | 0.405 | n.s. | -0.295 | 0.616 | n.s. | -0.289 | 0.614 | n.s. | 0.322 | 0.374 | n.s. | 0.012 | 0.496 | n.s. |
| i8 | 1.128 | 0.130 | n.s. | -0.994 | 0.840 | n.s. | 0.106 | 0.458 | n.s. | 0.773 | 0.220 | n.s. | -0.275 | 0.609 | n.s. |
| i9 | 0.711 | 0.239 | n.s. | -0.445 | 0.672 | n.s. | 0.554 | 0.290 | n.s. | 0.734 | 0.231 | n.s. | -0.130 | 0.552 | n.s. |
| i10 | 2.267 | 0.012 | 0.562 | 6.464 | $5.108 \times 10^{-11}$ | 0.450 | 9.011 | $\sim 0.0$ | 0.497 | 14.015 | $\sim 0.0$ | 0.497 | 20.114 | $\sim 0.0$ | 0.508 |

n.s. = not significant.

### Validating Population-Level Hybridization Detection

To contrast with the previous simulation, and to validate hybridization at the population level, we repeated the same procedure as above for simulating DNA sequence data for four taxa, but this time *all* individuals in the `sp2` population were 10% admixed. This maintains the overall level of 10% admixture in the population, but now it is distributed evenly among the individuals. We then ran a hybridization detection analysis on these simulated data using the *run_hyde.py, bootstrap_hyde.py* (500 replicates), and *individual_hyde.py* scripts.

#### Results

At the population level, the analysis with *run_hyde.py* detected significant hybridization (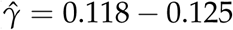) in all cases except for the simulation with 10 000 sites (Table 3). For this scenario, there was not enough data to detect the hybridization at the *α* = 0.05 level. Analyzing each individual in the `sp2` population correctly detected hybridization in all individuals, with estimates of 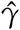 slightly over the original level of 10% admixture (Table 4). Bootstrap resampling of the individuals in the `sp2` population also demonstrated that hybridization was uniform, as can be seen by the unimodal distribution of 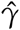 values across bootstrap replicates in Figure 3.

**Figure 3:**
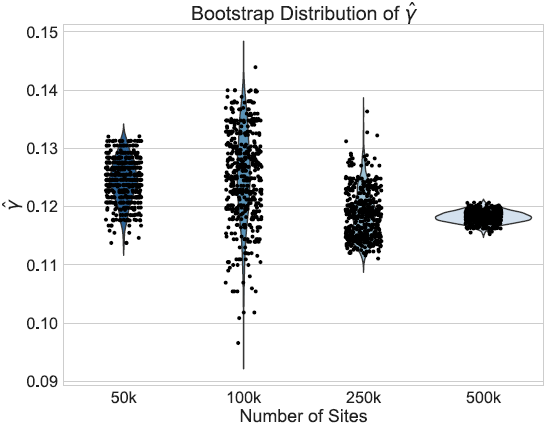
Violin plots of the distribution of 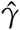 across 500 bootstrap replicates for 50 000, 100 000, 250 000, and 500 000 sites for the uniform hybridization simulations. The black dots in each violin plot represent the actual values of 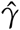 that were estimated by each replicate (black dots are jittered). The distribution of the estimates across bootstrap samples indicates that hybridization is uniform. The analyses with 10 000 sites did not have enough statistical power to detect hybridization and were therefore left out of the plot.

**Table 3:** Results of the population-level hybridization detection analyses for the 10% admixture simulation using the *run_hyde.py* script. Note that the result for the simulation with 10 000 sites was not significant at the *α* = 0.05 level (marked with *).

| # sites | Z-score | P-value | $\hat{\gamma}$ |
| --- | --- | --- | --- |
| 10 000* | 0.071 | 0.472 | 0.008 |
| 50 000 | 2.363 | 0.009 | 0.125 |
| 100 000 | 3.677 | $1.180 \times 10^{-4}$ | 0.125 |
| 250 000 | 5.796 | $3.401 \times 10^{-9}$ | 0.119 |
| 500 000 | 8.425 | $\sim 0.0$ | 0.118 |

**Table 4:**
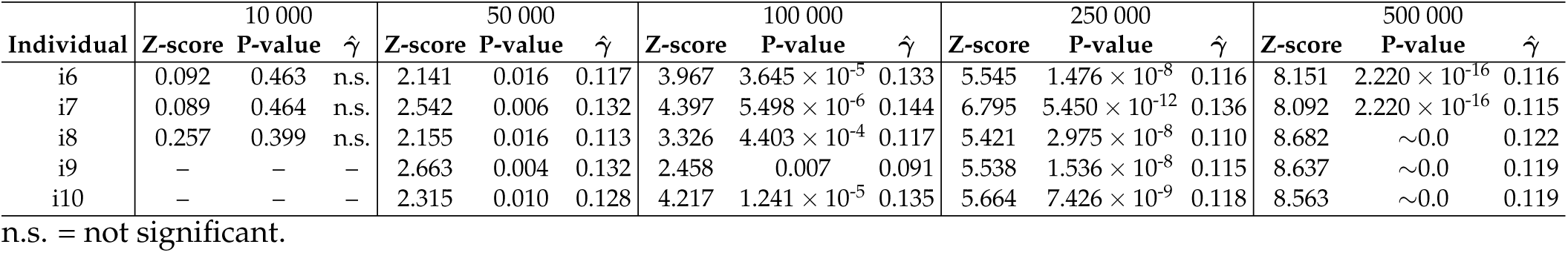
Results of the individual-level hybridization detection analyses for the 10% admixture simulation using the *individual_hyde.py* script. All individuals were equally admixed with *γ* = 0.1.

## Biological Examples

### Heliconius *Butterflies*

DNA sequence data from Martin et al. (2013) was downloaded for four populations of *Heliconius* butterflies (248 822 400 sites; Dryad: 10.5061/dryad.dk712). The number of individuals per population was as follows: four individuals of *H. melpomene rosina*, four individuals of *H. melpomene timareta*, four individuals of *H. cydno*, and one individual of *H. hecale*. We tested the hypothesis that *H. cydno* is a hybrid between *H. melpomene rosina* and *H. melpomene timareta* with *H. hecale* as an outgroup using the *run_hyde.py* script. We then conducted bootstrapping (500 replicates) of the *H. cydno* individuals, as well as testing each individual separately using the *bootstrap_ hyde.py* and *individual_hyde.py* scripts, respectively. As a last comparison, we reanalyzed the data with the *run_hyde.py* script, but changed the underlying code so that all missing or ambiguous sites were ignored in order to test the effect of including/excluding such data.

#### Results

Significant hybridization was detected in *H. cydno* at the population level (Z-score = 472.599, p-value = ~0.0, 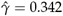). Each individual also showed significant levels of hybridization, with 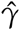 ranging from 0.324 to 0.393, indicating that hybridization in *H. cydno* is mostly uniform across the individuals sampled. Figure 4 shows a density plot for the estimated values of 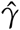 across the 500 bootstrap replicates. Although the distribution does exhibit some amount of jumping between different values they are still between the lower and upper bounds for the individual level estimates of 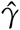, providing corroborating evidence that all the individuals in the *H. cydno* population are admixed. When we ignored missing or ambiguous sites, we still detected significant hybridization, but the value of the test statistic was less than half of the value from the original analysis (Z-score = 189.073, p-value = ~0.0, 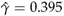). This demonstrates the increase in power that can result from including *all* relevant data in the analysis.

**Figure 4:**
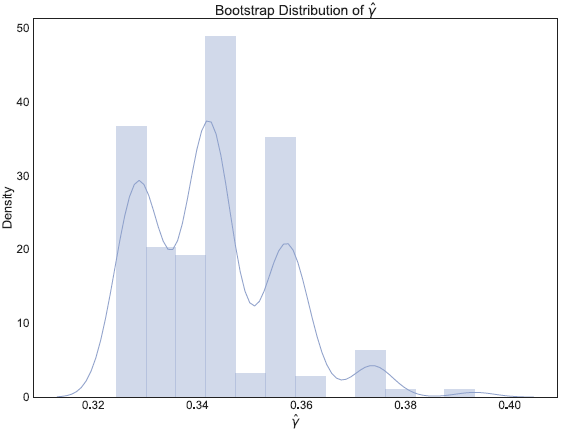
Density plot of the estimated values of 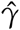 across 500 bootstrap replicates testing for hybridization in *Heliconius cydno* between *H. melpomene rosina* and *H. melpomene timareta*.

### *Swordtail Fish* (Xiphophorous)

Transcriptome data from Cui et al. (2013) was obtained from the authors for 26 species of swordtail fish (25 species of *Xiphophorous* + one outgroup from the genus *Priapella*). The data (3 706 285 sites) were first put in sequential Phylip format for analysis with HyDe. Each taxon was then analyzed at the individual level, resulting in 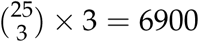 hypothesis tests. Because there were not multiple individuals per taxon, we only analyzed these data using the *run_hyde.py* script. As with the *Heliconius* example, we also reanalyzed these data ignoring all sites with missing or ambiguous bases.

#### Results

Out of 6900 tests, 2199 reported significant levels of hybridization (Bonferroni corrected p-value of 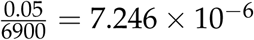). When ambiguous or missing bases were ignored, only 867 tests were reported as significant, again demonstrating the increase in power to detect hybridization when including these sites. The distribution of the 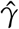 values for the 2199 significant tests is plotted in Figure 5. This plot shows that most of the admixture occurring in *Xiphophorous* is at low levels (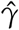 close to either 0.0 or 1.0). However, there are many instances of higher levels of admixture, which matches the findings of Cui et al. (2013) regarding the extensive amount of gene flow and introgression in this group. Solís-Lumis and Ané (2016) also analyzed these data and found evidence for an ancestral hybridization event between *X. xiphidium* and the northern swordtail clade (denoted NS in Solís-Lumis and Ané (2016), Fig. 10), a pattern we recover as well (259 significant hypothesis tests involving *X. xiphidium*).

**Figure 5:**
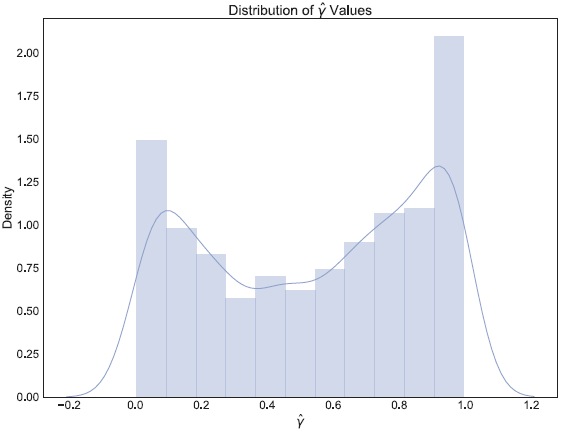
Density plot of the estimated values of 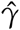 across the 2199 hypothesis tests that detected significant levels of hybridization in *Xiphophorous*.

### Timing

We timed all analyses for the *Heliconius* and swordtail data using the Unix `time` command. For *Heliconius*, the analysis using the *run_hyde.py* script took 11m 49.377s, and the analysis that ignored missing or ambiguous sites took 4m 47.667s. Analyzing the four individuals of *H. cydno* separately with the *individual_hyde.py* script took 10m 5.120s. Completing 500 bootstrap replicates for this data set ran the longest of any analysis that we conducted, taking a total of 4105m 33.312s (2.851 days). For the swordtail data, the analysis with the *run_hyde.py* script took 22m 40.673s, almost twice as long as the analysis for the *Heliconius* data. This indicates that analyzing more triples is more computationally intensive than analyzing more sites. When ambiguous or missing sites were ignored, the analysis took 5m 27.889s.

## Conclusions

Hybridization and gene flow are increasingly recognized as important forces in the evolution of taxa across the Tree of Life (Seehausen 2004; Arnold and Kunte 2017; Kagawa and Takimoto 2017). In addition, for groups such as plants, fish, and amphibians, hybridization can also be accompanied by whole genome duplication (allopolyploidy; Otto and Whitton 2000). Kamneva et al. (2017) recently used methods based on the coalescent with hybridization to model haplotypes in allopolyploid strawberries, and we anticipate that approaches using site pattern probabilities should be equally applicable in such situations. Gaining an understanding of when these complex processes are occurring and how they may vary among individuals within a population are important steps to understand the spatial distribution of shared genetic variation among diverging lineages. The methods we have implemented in HyDe to detect hybridization at the population and individual level provide a set of computationally efficient methods for researchers to assess patterns of admixture in natural populations. HyDe also provides an open source Python library for the calculation of site pattern probabilities that can easily be extended to calculate additional statistics that may be developed in the future. As access to genomic data continues to grow, we anticipate that methods such as HyDe that use phylogenetic invariants will play an important role for phylogenomic inferences in non-model species.

## Availability

HyDe is available as an open source Python package distributed under the GNU General Public License v3 for both Python 2.7 and 3.6. Developmental code can be found on GitHub (https://github.com/pblischak/HyDe) with official code releases uploaded to the Python Package Index (PyPI: https://pypi.python.org/pypi/phyde). Documentation for installing and running HyDe can be found on ReadTheDocs (http://hybridization-detection.readthedocs.io).

## Supplemental Materials

Simulated data sets, code, and other materials will be made available on the Dryad Digital Repository.

## Funding

This work was supported in part by grants from the National Science Foundation under the awards DMS-1106706 (JC and LSK) and DEB-1455399 (ADW and LSK), and National Institutes of Health Cancer Biology Training Grant T32-CA079448 at Wake Forest School of Medicine (JC).

## Acknowledgements

The authors thank Cécile Ané and Frank Burbink for the invitation to be a part of this special issue on the impact of gene flow and reticulation in phylogenetics, as well as David Posada and three anonymous reviewers for comments that improved the manuscript. We also thank R. Cui, C. Solís-Lemus, and C. Ané for their help with the *Xiphophorous* data set.

### Box 1

Example code to run hybridization detection analyses with HyDe using the Python API. The `phyde` module is loaded using the `import` command:

~~~
# import the phyde module
import phyde
~~~

Data are read in and stored using the *HydeData* class by passing the names of the input file, map file, and outgroup, as well as the number of individuals, the number of populations, and the number of sites.

~~~
# read in data using HydeData class
data = phyde.HydeData(“data.txt”, “map.txt”,
“out”, 16, 4, 50000)
~~~

With the data read in, we can use methods implemented in the *HydeData* class to test for hybridization at the population level [`test_triple(p1, hyb, p2)`], at the individual level [`test_individuals(p1, hyb, p2)`], and can bootstrap resample individuals [`bootstrap_triple(p1, hyb, p2, nreps)`].

~~~
# test for hybridization in the “sp2” population
res1 = data.test_triple(“sp1”, “sp2”, “sp3”)
# test all individuals in the “sp2” population
res2 = data.test_individuals(“sp1”, “sp2”, “sp3”)
# bootstrap individuals in the “sp2” population 200 times
res3 = data.bootstrap_triple(“sp1”, “sp2”, “sp3”, 200)
~~~

## References

Allman, E. S., L. S. Kubatko, and J. A. Rhodes. 2016. Split scores: a tool to quantify phylogenetic signal in genome-scale data. Systematic Biology 66:620–636.

Arnold, M. J. and K. Kunte. 2017. Adaptive genetic exchange: a tangled history of admixture and evolutionary innovation. Trends in Ecology and Evolution 32:601–611.

Baack, E. J. and L. H. Rieseberg. 2007. A genomic view of introgression and hybrid speciation. Current Opinions in Genetics and Development 17:513–518.

Behnel, S., R. Bradshaw, C. Citro, L. Dalcin, D. Seljebotn, and K. Smith. 2010. Cython: the best of both worlds. Computing in Science Engineering 13:31–39.

Cavender, J. A. and J. Felsenstein. 1987. Invariants of phylogenies in a simple case with discrete states. Journal of Classification 4:57–71.

Chifman, J. and L. S. Kubatko. 2014. Quartet inference from SNP data under the coalescent model. Bioinformatics 30:3317–3324.

Chifman, J. and L. S. Kubatko. 2015. Identifiability of the unrooted species tree topology under the coalescent model with time-reversible substitution processes, site-specific rate variation, and invariable sites. Journal of Theoretical Biology 374:35–47.

Cui, R., M. Schumer, K. Kruesi, R. Walter, P. Andolfatto, and G. G. Rosenthal. 2013. Phylogenomics reveals extensive reticulate evolution in Xiphophorus fishes. Evolution 67:2166–2179.

Durand, E. Y., N. Patterson, D. Reich, and M. Slatkin. 2011. Testing for ancient admixture between closely related populations. Molecular Biology and Evolution 28:2239–2252.

Eaton, D. A. R. and R. H. Ree. 2013. Inferring phylogeny and introgression using RADseq data: an example from the flowering plants (Pedicularis: Orobanchaceae). Systematic Biology 62:689–706.

Gerard, D., H. L. Gibbs, and L. S. Kubatko. 2011. Estimating hybridization in the presence of coalescence using phylogenetic intraspecific sampling. BMC Evolutionary Biology 11:291.

Green, R. E., J. Krause, A. W. Briggs, T. Maricic, U. Stenzel, M. Kircher, N. Patterson, H. Li, W. Zhai, M. H.-Y Fritz, N. F. Hansen, E. Y. Durand, A.-S Malaspinas, J. D. Jensen, T. Marques-Bonet, C. Alkan, K. Prüfer, M. Meyer, H. A. Burbano, J. M. Good, R. Schultz, A. Aximu-Petri, A. Butthof, B. Höber, B. Höffner, M. Siegemund, A. Weihmann, C. Nusbaum, E. S. Lander, C. Russ, N. Novod, J. Affourtit, M. Egholm, C. Verna, P. Rudan, D. Brajkovic, Ž. Kucan, I. Gušic, V. B. Doronichev, L. V. Golovanova, C. Lalueza-Fox, M. de la Rasilla, J. Fortea, A. Rosas, R. W. Schmitz, P. L. F. Johnson, E. E. Eichler, D. Falush, E. Birney, J. C. Mullikin, M. Slatkin, R. Nielsen, J. Kelso, M. Lachmann, D. Reich, and S. Pääbo. 2010. A draft sequence of the Neandertal genome. Science 328:710–722.

Hasegawa, M., H. Kishino, and T. Yano. 1985. Dating of human-ape splitting by a molecular clock of mitochondrial DNA. Journal of Molecular Evolution 22:160–174.

Hudson, R. R. 2002. Generating samples under a Wright-Fisher neutral model of genetic variation. Bioinformatics 18:337–338.

Joly, S., P. A. McLenachan, and P. J. Lockhart. 2009. A statistical approach for distinguishing hybridization and incomplete lineage sorting. American Naturalist 174:E54–E70.

Jukes, T. H. and C. R. Cantor. 1969. Evolution of protein molecules. Pages 21–132, in Mammalian protein metabolism (H. Monroe, ed.). New York: Academic Press.

Kagawa, K. and G. Takimoto. 2017. Hybridization can promote adaptive radiation by means of transgressive segregation. Ecology Letters 21:264–274.

Kamneva, O. K., J. Syring, A. Liston, and N. A. Rosenberg. 2017. Evaluating allopolyploid origins in strawberries (fragaria) using haplotypes generated from target capture sequencing. BMC Evolutionary Biology 17:180.

Kingman, J. F. C. 1982. On the genealogy of large populations. Journal of Applied Probability 19:27–43.

Kubatko, L. S. 2009. Identifying hybridization events in the presence of coalescence via model selection. Systematic Biology 58:478–488.

Kubatko, L. S. and J. Chifman. 2015. An invariants-based method for hybridization detection from genome-scale sequence data. bioRxiv doi:10.1101/034348.

Lake, J. A. 1987. A rate-independent technique for analysis of nucleic acid sequences: evolutionary parsimony. Molecular Biology and Evolution 4:167–191.

Maddison, W. P. 1997. Gene trees in species trees. Systematic Biology 46:523–536.

Mallet, J. 2005. Hybridization as an invasion of the genome. Trends in Ecology and Evolution 20:229–237.

Mallet, J. 2007. Hybrid speciation. Nature 446:279–283.

Martin, S. H., K. K. Dasmahapatra, N. J. Nadeau, C. Salazar, J. R. Walters, F. Simpson, M. Blaxter, A. Manica, J. Mallet, and C. D. Jiggins. 2013. Genome-wide evidence for speciation with gene flow in Heliconius butterflies. Genome Research 23:1817–1828.

Martin, S. H., J. W. Davey, and C. D. Jiggins. 2014. Evaluating the use of ABBA-BABA statistics to locate introgressed loci. Molecular Biology and Evolution 32:244–257.

Meng, C. and L. S. Kubatko. 2009. Detecting hybrid speciation in the presence of incomplete lineage sorting using gene tree incongruence: a model. Theoretical Population Biology 75:35–45.

Otto, S. P. and J. Whitton. 2000. Polyploid incidence and evolution. Annual Review of Genetics 34:401–437.

Patterson, N., P. Moorjani, Y. Luo, M. Swapan, N. Rohland, Y. Zhan, T. Genschoreck, T. Webster, and D. Reich. 2012. Ancient admixture in human history. Genetics 192:1065–1093.

Pease, J. B. and M. W. Hahn. 2015. Detection and polarization of introgression in a five-taxon phylogeny. Systematic Biology 64:651–662.

Rambaut, A. and N. C. Grassly. 1997. Seq-Gen: an application for the Monte Carlo simulation of DNA sequence evolution along phylogenetic trees. Computer Applications in the Biosciences 13:235–238.

Seehausen, O. 2004. Hybridization and adaptive radiation. Trends in Ecology and Evolution 19:198–207.

Solís-Lumis, C. and C. Ané. 2016. Inferring phylogenetic networks with maximum pseudolikelihood under incomplete lineage sorting. PLoS Genetics 12:e1005896.

Tavaré, S. 1986. Some probabilistic and statistical problems in the analysis of DNA sequences. Lectures on Mathematics in the Life Sciences 17:57–86.

Tian, Y. and L. S. Kubatko. 2017. Rooting phylogenetic trees under the coalescent model using site pattern probabilities. BMC Evolutionary Biology 17:263.

Yu, Y., R. M. Barnett, and L. Nakhleh. 2013. Parsimonious inference of hybridization in the presence of incomplete lineage sorting. Systematic Biology 62:738–751.

Yu, Y., J. H. Degnan, and L. Nakhleh. 2012. The probability of a gene tree topology within a phylogenetic network with applications to hybridization detection. PLoS Genetics 8:e1002660.

Yu, Y., J. Dong, K. J. Liu, and L. Nakhleh. 2014. Maximum likelihood inference of reticulate evolutionary histories. Proceedings of the National Academy of Sciences 111:16448–16453.

Yu, Y., C. Than, J. H. Degnan, and L. Nakhleh. 2011. Coalescent histories on phylogenetic networks and detection of hybridization despite incomplete lineage sorting. Systematic Biology 60:138–149.

